# Out-of-Register Aβ_42_ Assemblies as Models for Neurotoxic Oligomers and Fibrils

**DOI:** 10.1101/190835

**Authors:** Wenhui Xi, Elliott K. Vanderford, Ulrich H.E. Hansmann

## Abstract

We propose a variant of the recently found S-shaped Aβ_1‒42_-motif that is characterized by out-of-register C-terminal β-strands. We show that chains with this structure can not only form fibrils that are compatible with the NMR signals, but also barrel-shaped oligomers that resemble the ones formed by the much smaller cylindrin peptides. Running at physiological temperatures long all-atom molecular dynamics simulations with an explicit solvent, we study the stability of these constructs and show that they are plausible models for neurotoxic oligomers. Analyzing the transitions between different assemblies we suggest a mechanism for amyloid formation in Alzheimer’s disease.

## Introduction

Alzheimer’s disease and an increasing number of other illnesses are connected with the presence of amyloid fibrils.^1-4^ The bottleneck in the emergence of mature fibrils is the time needed for self-assembly of monomers into a critical nucleus^5^. In this process, smaller oligomers are formed that in the case of Alzheimer’s disease appear to be the main neurotoxic agents causing memory loss^6,7^. It is believed that these oligomers, made of 37-42 residue long Aβ peptides, damage cells by disrupting cell membranes, and that the cell toxicity and disease stage are related to the different conformation of Aβ oligomer or fibrils.^8,9^ However, the exact mechanism of neurotoxicity is not clear^10,11^

This is in part because it is a challenge to resolve experimentally the structure of such oligomers, existing in an ensemble of transient and polymorphic forms^12-15^. One of the few experimental models is cylindrin, the eleven-residue segment K11V of αB – crystalline, which in solution can assemble not only into fibrils with the typical in-register β-strands but also into fibril-like structures with out-of-register β-strands,^16^ and, most interesting, into a barrel-shaped hexamer, that is characterized by out-of-register antiparallel β-strands arranged as a cylindrical barrel and hold together by inter-chain hydrogen bonds. The same barrel motif can also be formed by three tandem-repeats K11V-TR, each built out of two K11V segments linked by two glycine residues, where the tandem repeats fold into a U-shaped conformation.

While the Eisenberg group has proposed that the cylindrin barrel motif is shared by the various toxic oligomers in amyloid diseases,^17^ its actual relevance for the pathogenesis of Alzheimer’s disease is not clear. Thanh D. et al^18^ have proposed cylindrin-like models of tandem repeats of eleven-residue-long Aβ-fragments (residue 24-34, 25-35, 26-36), hold together by two glycine residues; and have studied the stability of these models with a combination of computational modeling and ion-mobility mass spectrometry, electron microscopy and other experimental techniques. However, unlike for such short fragments, it is difficult to form cylindrin-like barrels, made of out-of-register antiparallel β-strands, from *full-sized* Aβ-peptides if one assumes for the individual chains the U-shaped structure seen in most of the Aβ-fibril models in the Protein Data Bank.^8,12-15^ This is because the phenylalanine rings of residues 19 and 20 do not fit into the core of a β- barrel made of three or four U-shaped chains. The steric constraint disappears for larger oligomers, which, however, will have an unstable core.

It was recently shown that, unlike Aβ_1‒40_ peptides, the less common but more toxic Aβ_1‒42_ species can form fibrils made from three-stranded S-shaped chains. Assemblies of Aβ_1‒42_ chains taking this fold are not restricted by the above described steric constraints.^19^ As Aβ_1‒40_ cannot assume the S-shape motif^19^, one can speculate that the higher toxicity of Aβ_1‒42_ is related to its ability to assemble into cylindrin-like barrel-shaped oligomers.

As a first step into testing this hypothesis we will present in this article a variant of the S-shaped Aβ_1‒42_-motif with out-of-register C-terminal β-strand that similar to cylindrin can assemble into both fibrils and barrel-shaped oligomers. The fibril assemblies are compatible with the NMR signals. We show that our constructs are possible models for neurotoxic oligomers and study their stability at physiological temperatures through long all-atom molecular dynamics simulations with an explicit solvent. Analyzing these trajectories, we propose a mechanism for amyloid formation in Alzheimer’s disease.

## Methods

### Model construction

While in all known wild type Aβ_1-40_ amyloid fibrils the individual chains form two β-strands connected by a loop region,^12-15^ several recent studies have demonstrated the possibility of other chain arrangements for Aβ_1-42_ fibrils.^19-23^ These fibril models are characterized by S-shaped chains, with the N-terminal strand β1 made of residues 12‒18, the central strand β2 of residues 24‒33, and the C-terminal strand β3 of residues 36‒40.^19^ We have shown in earlier work that this S-shaped conformation is not stable in assemblies of Aβ_1‒40_ peptides.^24^ This is not because the lack of the intra-chain salt bridge between side chain of residue K28 and the main chain of residue A42, that connects the β2 and β3 strands and cannot be formed in Aβ_1‒40_ peptides, but because of the lack of hydrophobic contacts involving the C-terminal residues I41 and A42. As already observed in Ref. ^24^, the β2-turn-β3 motif is surprisingly stable. For instance, the hydrogen bonding in this region can change from *one between different chains* to one between the *strands of the same chain*, with only a small decrease in the total number of hydrogen bonds. In a U-shape fibril model, the loss of inter-chain hydrogen bonds would dissolve the fibril; however, with a S-shaped geometry the fibril will be still stabilized by inter-chain hydrogen bonds connecting the β1-strands. It is this property that allows us to construct our models.

In order to build an out-of-register fibril model that preserves as much as possible the interchain contacts of the in-register fibril, we start with the triple β-stranded Aβ42 fibril model (PDB id: 2MXU) resolved by Xiao et al^19^, which we call in this paper the In-Register-Fibril (IRF) model and show in Figure 1a. The IRF model is built from three-stranded S-shaped chains hold together by *inter-chain* hydrogen bonds between neighboring chains. Using AMBERTOOLS^25^ we alter in each chain the β2-turn-β3 segment into a β-turn between residue 27 to 42 while keeping the salt bridge K28-A42, but replacing the inter-chain hydrogen bonds by out-of-register *intra-chain* hydrogen bonds. We tried different possible staggering patterns by minimizing them and testing their stability in molecular dynamics simulations of 20 ns. In this way, we derived an out-of-register structure with the hydrogen bond staggering that is listed in table 1. We then assemble six copies of this structure into a fibril-like hexamer that is kept together by inter-chain hydrogen bonds between β1 strands of neighbouring chains. Minimizing and equilibrating by 50 ns molecular dynamics runs led to four models of out-of-register fibrils, out of which we choose the most stable one as our candidate structure for an Out-of-Register Fibril (ORF) model shown in Figure 1b. The in-register part, residues 11-26, is colored in blue and the out-of-register sections are shown in yellow.

**Table 1.** Hydrogen-bonding between backbone atoms in the core region (residues 27 to 42) of the out-of-register fibril (ORF) and the out-of-register β-barrel models BB3 and BB4. All models share the same intra-chain hydrogen-bonds, but inter-chain hydrogen bonds exist in this region only in the barrels BB3 and BB4.

| Intra-chain hydrogen-bonds |  | Inter-chain hydrogen-bonds (Only in BB3 and BB4) |  |  |  |
| --- | --- | --- | --- | --- | --- |
|  |  | In-register |  | Out-of-register |  |
| β2 strand | β3 strand | Chain A | Chain B | Chain A | Chain B |
| 40VAL:O | 29GLY:N | 28LYS :O | 34LEU :N | 31ILE :O | 39VAL :N |
| 40VAL:N | 29GLY:O | 30ALA :N | 32ILE :O | 31ILE :N | 39VAL :O |
| 38GLY:O | 31ILE:N | 30ALA :O | 32ILE :N | 37GLY :O | 41ILE :N |
| 38GLY:N | 31ILE:O | 32ILE :N | 30ALA :O | 37GLY :N | 41ILE :O |
| 37GLY:N | 33GLY:O | 32ILE :O | 30ALA :N |  |  |
|  |  | 34LEU :O | 28LYS :O |  |  |

**Figure 1.**
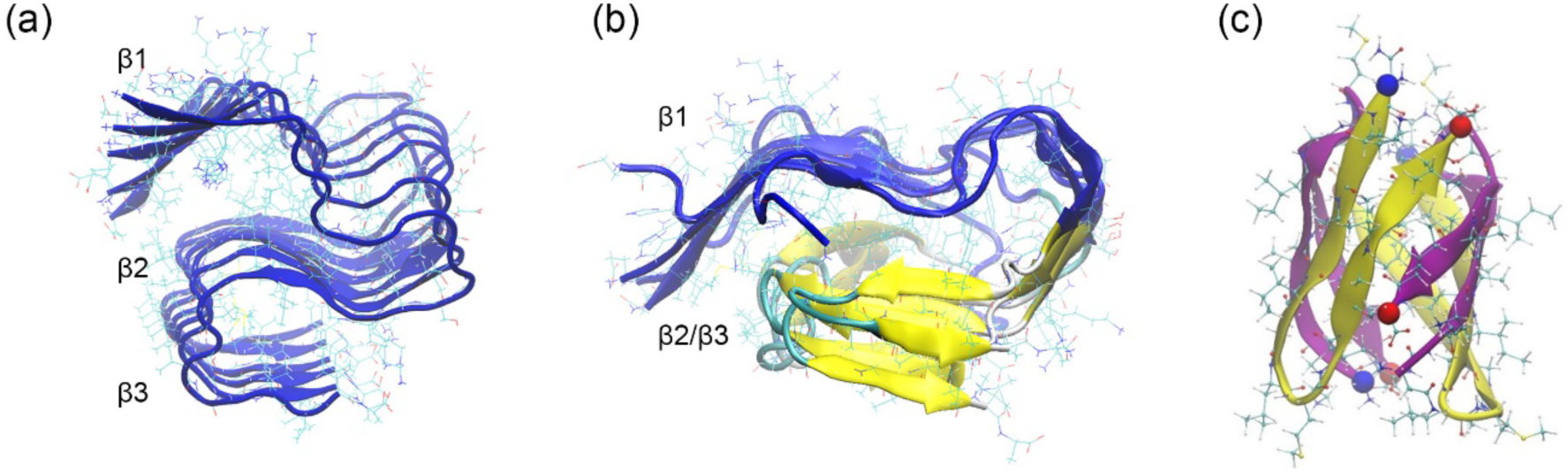
(a) Our start-configuration for the in-register (IRF) fibril model as derived from the experimentally determined structure (PDB-Id: 2MXU). (b) Representative conformation of our out-of-register (ORF) fibril model. The in-register section (residue 11-26) is drawn in blue, and the out-of-register parts in yellow. (c) Representative structure for our proposed β-barrel tetramer model. Neighboring chains are drawn alternating in yellow and purple. For each chain, either the β2-strand or the β3-strand forms in-register inter-chain hydrogen-bonds with one of its neighbor, while the other one will form out-of-register hydrogen-bonds with the other neighbor.

The same out-of-register Aβ_1‒42_ chain configuration that is utilized in the out-of-register fibril model is also used to build out-of-register β-barrel trimer (BB3) and tetramer (BB4) models. Different side chain orientations and backbone hydrogen bond pattern allow for a multitude of possible arrangements. We have built more than twelve different assemblies and chosen the ones with smallest fluctuation and lowest Generalized-Born energy. Our most stable tetramer is shown in Figure 1c, with adjacent chains drawn in in different color (yellow and purple), and the hydrogen bond patterns again listed in table 1. While the intra-chain hydrogen bond pattern is the same as in the ORF model, for each chain, either the β2 or the β3 strand forms in addition *in-register non-staggered inter-chain* hydrogen bonds with a neighbor chain, while the other strand forms *out-of-register inter-chain* hydrogen bonds with its neighbor.

Coordinates of the out-of-register fibril and barrel structures, generated by the above procedure, are available as supplemental material.

### Molecular dynamics set-up

Our models are constructed with the AmberTools tool suit^25^ and studied by all-atom molecular dynamics using the GROMACS 4.6.7 package^26^. These simulations rely on the CHARMM 36 force field^27^ and TIP3P solvent^28^. Restraining the bonds in protein and solvent with the LINCS^29^ and SETTLE^30^ algorithm allows us to set the integration time-step to 2 fs. A NPT ensemble is realized by setting the temperature to 310 K with by v-rescale thermostat^31^ and the pressure to 1 bar by a Parrinello-Rahman barostat^32^. In order to check the reliability of our results we have studied each model in two independent runs (starting from the same configuration but different initial velocity distributions that correspond to T=310K). For the out-of-register fibril model ORF each trajectory lasts for 200ns, while they last 100 ns for the trimer and tetramer β-barrel models BB3 and BB4.

The resulting trajectories are analyzed using the GROMACS tools suite. However, binding energies are approximated by MMGBSA,^33^ with the calculations done Amber14^25^ setting ‘igb=8’ ^34^ and omitting entropy terms. NMR signals for our proposed structures are predicted with SHIFTX2 approach^35^ using the web-sever *http://www.shiftx2.ca*.

### Replica Exchange with Tunneling

Transitions between in-register fibrils and out-of-register fibril involve crossing of high free-energy barriers that require impractically long simulation times when using regular molecular dynamics. In order to alleviate this problem we have recently introduced a new generalized-ensemble technique named replica-exchange-tunneling (RET) methods.^36^ Unlike in temperature-driven replica-exchange sampling, the acceptance probability is not calculated at the time of the exchange move but only after the exchanged replicas have evolved over a short micro-canonical trajectory. By this procedure one can “tunnel” through unfavorable “transition” states resulting from the exchange move. For details of the sampling technique, see references^36,37^.

In the present paper, we use this improved exchange move to enhance conversions between out-of-register and in-register fibrils by building a chain of replicas that differ in how strongly they are biased to either of the two models. For this purpose we define an energy function E_pot_ = E_phys_+E_Go_ + λ * E_λ_, where E_phy_ describes the energy of the “physical” model, E_Go_ the energy of a suitable Go-model biasing toward either the IRG or the ORG model, and E_λ_ describes the similarity of configurations in physical and Go-model. The strength of coupling between the two models is set by the parameter λ and varies between the replicas. Note that the exchange move depends only on the coupling term, i.e., is proportional to

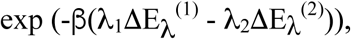

when two configurations are exchanged between replicas with coupling parameters λ_1_ and λ_2_. A consequence of this procedure is a walk of replicas through the space of coupling parameters, with at one end maximal bias toward the in-register-fibril structure IRF, and at the opposite end maximal bias toward the out-of-register fibril ORF. In its most straightforward implementation of this approach are data only analyzed for the replica where λ = 0, i.e., where the physical model is not biased by other terms. However, information from replicas with λ ≠ 0 can be also utilized by using re-weighting techniques^37^. More details on this approach can be found in previous work where we have demonstrated the superior sampling of RET in studies of switching proteins^36^ and studies of conformational changes in small oligomers^37^.

Our RET algorithm is implemented in a modified version Gromacs 4.6.5 that is available on request from the authors. In order to speed up our simulations we rely on a combination of the CHARMM36 force field ^27^ with a generalized-Born model^38^ to approximate protein-solvent interaction by an implicit solvent. The Go-interaction terms are given by the SMOG energy function as generated by the SOMG-server^39^ at http://smog-server.org. We use 32 replicas with λ = (0.015, 0.0135, 0.012, 0.01, 0.008, 0.006, 0.005, 0.004, 0.003, 0.002, 0.001, 0.00075, 0.0005, 0.0003, 0.0001, 0, 0, 0.0001, 0.0003, 0.0005, 0.00075, 0.001, 0.002, 0.003, 0.004, 0.005, 0.006, 0.008, 0.01, 0.012, 0.0135, 0.015). Our implementation does not allow one to have all replicas the same temperature, instead the thermostat temperature changes in steps of 0.1K between 330K and 330.31K. All the replicas start from the same initial conformations obtained as the final configurations of a 20 ns long molecular dynamics run at 500K with the β1-strand restrained, starting from the IRF model (see Figure 1a) and leading to disordered β-2 and β-3 sections. The integration step of RET run is 2 fs and exchange attempt are made every 20ps. Each replica trajectory lasted for 50ns, but only the last 25ns are used for analysis. As in previous work^37^ we analyze the energy landscape of the fibril-like hexamer by reweighting data from all replica to the case of λ=0, i.e., the case where the “physical” model is not biased by the Go-term.

## Results and Discussion

### An out-of-register fibril model of Aβ42

Changing the *inter-chain* hydrogen bonds, connecting the β-strands of one chain with the corresponding strands of the next chain, to *intra-chain* hydrogen bonds, connecting the two strands into a β-bend, will dissolve a fibril built out of U-shaped Aβ40 or Aβ42 chains. This is different for the triple-stranded S-shaped Aβ42 fibrils where *inter-chain* hydrogen bonds, connecting the β2 and β3 strands of one chain with the corresponding stands in the next chain, can be replaced by *intra-chain* hydrogen bonds connecting the two strands in each chain into a β-bend. As the chains are still hold together by *inter-chain* hydrogen bonds connecting the β1 strands, it is not a priori necessary that our ORF model is unstable and must decay. For this reason, we start our investigation by analyzing first the actual stability of our out-of-register fibril ORF model. For this purpose, we have followed the time evolution of this fibril fragment in two independent molecular dynamics simulations of 200ns, with the simulation protocol described in the method section.

Visual inspection of the trajectories shows that during the simulations, the ORF hexamer remains stable, however, not to the extent the IRF hexamers (PDB id: 2MXU) are preserved in corresponding simulations. This observation can be quantified by measuring in both cases the root-mean-square-deviation (RMSD) to the respective start configurations along the two trajectories for each system. In Figure 2(a) we show this quantity as function of time for both ORF and IRF fibril fragments. Displayed are for each system the values from the trajectory where the final RMSD value was largest. In both models, neither the β1-strand nor the β2-β3 bend are dissolved at the end of 200ns simulations; and while the RMSD values are higher in the ORF case than for the IRF model, measurements of the root-mean-square fluctuations (RMSF) show that the β2-β3 bend is only slightly more flexible in the out-of-register fibril than in the in-register fibril, see Figure 2(b). Note that the turn region itself is even less flexible in the ORF model than in the IRF fibril.

**Figure 2.**
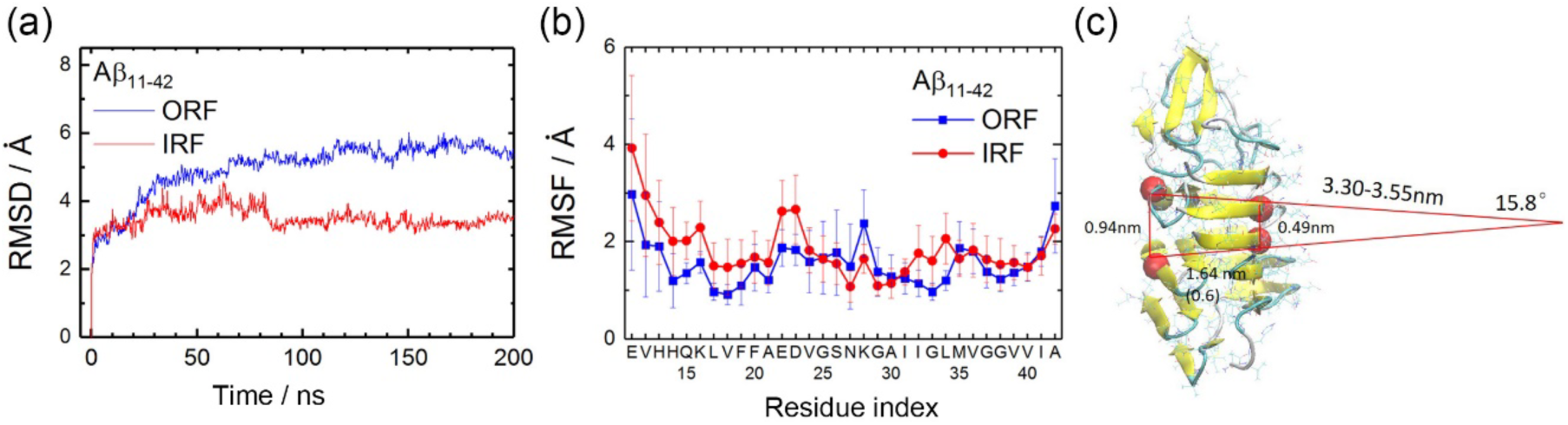
(a) The root-mean-square-deviation (RMSD) with respect to the start configurations as function of time measured in our simulations of Aβ_11-42_ hexamer, either assembled as in-register IRF fibril model (red) or as the out-of-register ORF fibril model (blue). (b) The root-mean-square-fluctuations (RMSF) of each residue measured for both assemblies. (c) Radius of curvature and bend angle per chain in the out-of-register fibril ORF after a molecular dynamics run of 200 ns. The Cα-atoms of residues V17 and V40 are drawn in red.

In our out-of-register fibril model ORF, the first intra-chain hydrogen bond connecting the β2 and β3 strands is formed between nitrogen atom of residue G29 and the oxygen atom of residue V40, with the other hydrogen bonds formed accordingly (the final one connecting residue 33 and residue 37). Although the hydrogen bonds connect in the initial configuration β2 and β3 regions that are arranged perpendicular to the direction of fibril elongation, after 200ns the hydrogen bonds have rotated in the final configuration by about 20-degree leading to a looser side-chain packing in the β2 region. On the other hand, the β1-strands stay unchanged in both models. However, the transition from inter-chain into intra-chain hydrogen bonds leads to a larger average distance between β2 and β3 regions of adjacent chains than seen between the N-terminal β1-region, see Figure 2(c). Hence, the ORF fibril is after thermalizing not straight but bend by an angle of about 15.8° per chain. This bending angle would lead to a tire-shaped fibril with an outer radius of ~5.1 nm, consisting of 22-23 chains. In the out-of-register section (the β2 and β3 hairpin), inter-chain contacts are not by hydrogen bonds between main chain atoms but are between side-chains. Hence, the residues in this section, for example, Leu or Val, are not as tightly packed than in the in-register form, which consequently leads to the above mentioned larger average inter-chain distances in this part but not seen between the β1-strands. Measuring the distance between residues V17 and V40 (shown as red balls in Figure 3) we estimate the radius of curvature as about 3.4nm. In a hybrid fibril where ORF and IRF assemblies would alternate, such twist of the fibril could lead to the “sharp turns” widely observed in Transmission Electron Microscopy or Atomic Force Microscopy images of Aβ fibrils. Note that in pure in-register-fibrils such sharp turns or twists, while in principle possible, require breaking of hydrogen bonds and are therefore energetically costly.

**Figure 3.**
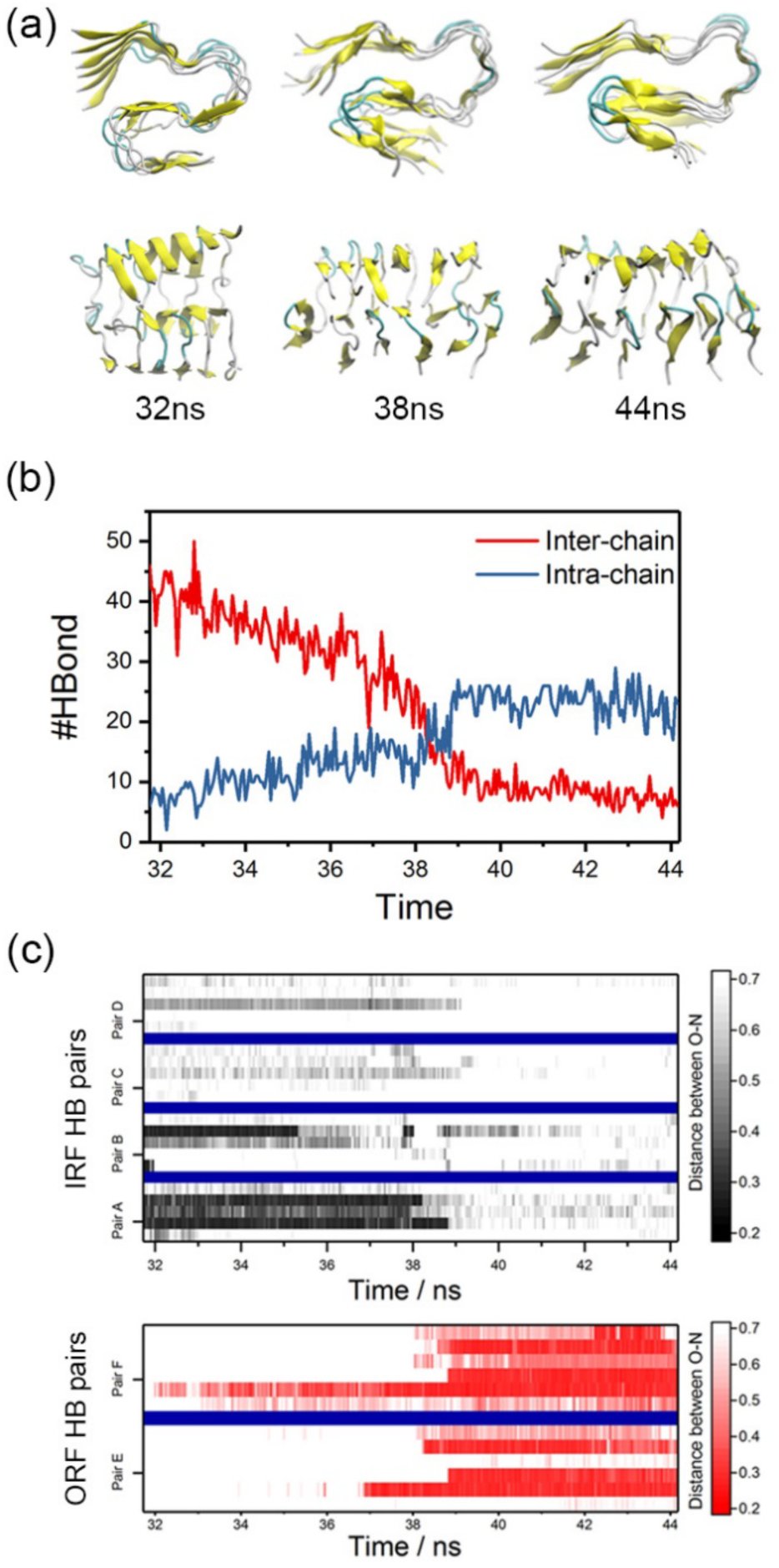
Representative configurations of a Aβ_1-42_ hexamer walking between a replica with strong bias toward the in-register fibril model IRF and a replica where the physical model is biased toward the out-of-register model ORF. Note that the inter-chain hydrogen bonds in the β2-turn-β3 region gradually dissolve and are replaced by intra-chain hydrogen bonds. This is quantified in (b) where we show along this tunneling event the number of inter-chain and intrachain hydrogen bonds involving residues in the C-terminal β2-turn-β3 motif; and in (c) where this time evolution is shown in more detail for the four pairs of inter-chain hydrogen bonds seen in the IRF fibril (A and B connecting β2-strands, C and D β3-strands) and the two pairs of intrachain hydrogen bonds seen in the ORF fibril (E connecting residues 29 and 40, F residues 33 and 37).

The in-register fibril model IRF has been derived from ss-NMR (solid-state Nuclear Magnetic Resonance) measurements.^19^ However, the relation between NMR signal and molecule structure is not straightforward. Different structures can be within the error bars compatible with a given set of measurements. For this reason, we have compared our out-of-register model ORF with the NMR signals that led to the in-register fibril structure as deposited in the PDB under identifier 2MXU. For this purpose, we have predicted the ^13^C and ^15^N chemical shifts that would result from our structure, using the SHIFTX2 software^35^ with the coordinates of our model as input. The so-calculated shifts are shown in Table 2. Note that the experimental chemical shift of some residues, such as E22, N27 and M35, are not given in Ref.^19^, and therefore are also not listed in Table 2. Interestingly, most of the predicted shifts differ by less than 10% from the experimental measurements, and for the backbone are the differences between the two models in the signals of CO and NH atoms around 2% to 5%. These small differences suggest that our model is consistent with the experimental measured chemical shifts, even for the out-of-register β2 and β3 regions, where the two fibril models differ most. Hence, the NMR data do not exclude our above-mentioned hypothesis of Aβ42 fibrils having a hybrid organization, with most parts being in-register assemblies, but parts of it having an out-of-register hydrogen bonding that leads to the experimentally observed twists in the fibril architecture.

**Table 2.**
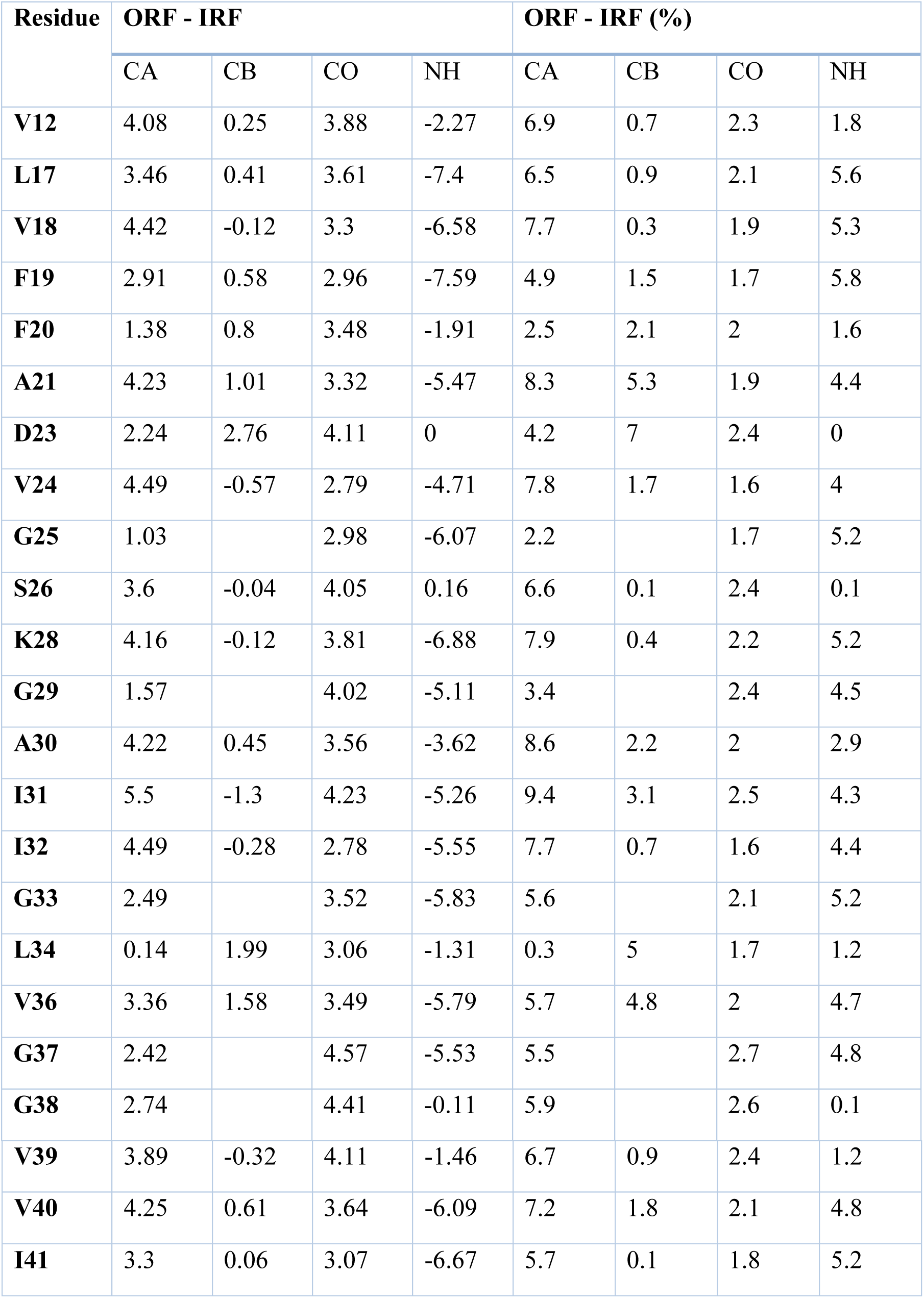
Comparison of ^13^C and ^15^N chemical shifts as predicted for our out-of-register fibril model ORF with the experimental results in previous work by Y. Xiao et al^19^ to propose the in-register fibril model IRF (PDB-ID 2MXU).

| Residue | ORF - IRF |  |  |  | ORF - IRF (%) |  |  |  |
| --- | --- | --- | --- | --- | --- | --- | --- | --- |
|  | CA | CB | CO | NH | CA | CB | CO | NH |
| **V12** | 4.08 | 0.25 | 3.88 | -2.27 | 6.9 | 0.7 | 2.3 | 1.8 |
| **L17** | 3.46 | 0.41 | 3.61 | -7.4 | 6.5 | 0.9 | 2.1 | 5.6 |
| **V18** | 4.42 | -0.12 | 3.3 | -6.58 | 7.7 | 0.3 | 1.9 | 5.3 |
| **F19** | 2.91 | 0.58 | 2.96 | -7.59 | 4.9 | 1.5 | 1.7 | 5.8 |
| **F20** | 1.38 | 0.8 | 3.48 | -1.91 | 2.5 | 2.1 | 2 | 1.6 |
| **A21** | 4.23 | 1.01 | 3.32 | -5.47 | 8.3 | 5.3 | 1.9 | 4.4 |
| **D23** | 2.24 | 2.76 | 4.11 | 0 | 4.2 | 7 | 2.4 | 0 |
| **V24** | 4.49 | -0.57 | 2.79 | -4.71 | 7.8 | 1.7 | 1.6 | 4 |
| **G25** | 1.03 |  | 2.98 | -6.07 | 2.2 |  | 1.7 | 5.2 |
| **S26** | 3.6 | -0.04 | 4.05 | 0.16 | 6.6 | 0.1 | 2.4 | 0.1 |
| **K28** | 4.16 | -0.12 | 3.81 | -6.88 | 7.9 | 0.4 | 2.2 | 5.2 |
| **G29** | 1.57 |  | 4.02 | -5.11 | 3.4 |  | 2.4 | 4.5 |
| **A30** | 4.22 | 0.45 | 3.56 | -3.62 | 8.6 | 2.2 | 2 | 2.9 |
| **I31** | 5.5 | -1.3 | 4.23 | -5.26 | 9.4 | 3.1 | 2.5 | 4.3 |
| **I32** | 4.49 | -0.28 | 2.78 | -5.55 | 7.7 | 0.7 | 1.6 | 4.4 |
| **G33** | 2.49 |  | 3.52 | -5.83 | 5.6 |  | 2.1 | 5.2 |
| **L34** | 0.14 | 1.99 | 3.06 | -1.31 | 0.3 | 5 | 1.7 | 1.2 |
| **V36** | 3.36 | 1.58 | 3.49 | -5.79 | 5.7 | 4.8 | 2 | 4.7 |
| **G37** | 2.42 |  | 4.57 | -5.53 | 5.5 |  | 2.7 | 4.8 |
| **G38** | 2.74 |  | 4.41 | -0.11 | 5.9 |  | 2.6 | 0.1 |
| **V39** | 3.89 | -0.32 | 4.11 | -1.46 | 6.7 | 0.9 | 2.4 | 1.2 |
| **V40** | 4.25 | 0.61 | 3.64 | -6.09 | 7.2 | 1.8 | 2.1 | 4.8 |
| **I41** | 3.3 | 0.06 | 3.07 | -6.67 | 5.7 | 0.1 | 1.8 | 5.2 |

### Transitions between in-of-register and out-register fibril organizations

Our above conjecture assumes that at least locally S-shaped Aβ_1-42_-chains can alter their hydrogen bonding from an in-register fibril organization toward an out-of-register fibril, and vice versa. As the free-energy barrier between the two fibril organizations could be high, such transitions may be difficult to observe in regular all-atom molecular dynamics. For this reason, we use Replica-Exchange with Tunneling (RET), a sampling technique developed specifically to cross such high energy barriers, to study the transition between IRF and ORF fragments. Our simulation protocol is described in the Method Section and the resulting simulations realize a random walk between replicas where the physical model is biased strongly toward the in-register-fibril organization of the IRF model, and, on the other side, replicas where the physical model is biased toward the out-of-register organization of the ORF model. In Figure 3 we show three representative configurations sampled along one such walk that started at the IRF model and led to the out-of-register ORF model. For this walk we also show in Figure 3(b) and (c) the change in hydrogen bonding as observed between the third and fourth chain. As the Aβ_1‒42_ hexamer converts from an in-register fibril organization to the out-of-register organization, the inter-chain hydrogen bonds in the β2-turn-β3 region gradually dissolve, starting in the turn region, and at a later stage are replaced by intra-chain hydrogen bonds, again formed first close to the turn. This transition starts from surface chains and propagates inside the fibril fragment. The smoothness of this transition implies that the our-of-register fibril could be an on-pathway intermediate for Aβ42 fibrils.

Note, however, that because of the artificial dynamics in RET, the above described walk is not necessarily the most likely one. For this being the case, the walk has to be consistent with the free-energy (potential of man force) landscape of the system, a quantity widely monitored in computational studies of amyloid peptide^40,41^. We show this landscape in Figure 4 projected on the number of inter-chain and intra-chain hydrogen bonds involving residues in the β2-turn-β3 turn. The IRF model here has only inter-chain hydrogen bonds, and the ORF fibril only intrachain hydrogen bonds. The free energies are obtained by re-weighting data from all 32 replicas onto the case of λ=0, i.e., for the case where the physical model is not biased toward either the IRF or the ORF model. The local minimum corresponding to the IRF-fibril (marked as (a) in the figure) is chosen as zero in the free-energy landscape. Going from the IRF form to the ORF form requires to cross a free energy barrier of 3.1 RT, where the transition state (d) is characterized by configurations in which the inter-chain hydrogen bonds connecting β3-strands between neighboring chains are dissolved but not the hydrogen bonds that connect the β2-strands. Disintegration of these hydrogen bonds lowers again the free energy and leads to the intermediate state (c) which has a relative free energy of 1.7 RT where only about 10% of the segments made of residue 27 to 32 (the β2-strand region) maintain inter-chain hydrogen bonds, and most of the residues in turn (residue 33-36) and β3 section (residues 37-42) have a random coil or extended structure. The rearrangements needed for formation of intra-chain hydrogen bonds between the β2 and β3 strands of the various chains initially raises the free energy by about 1 RT before the gain in energy by forming the hydrogen bonding of the ORF fibril (marked (b) in the figure) lowers the free energy again to a value only 0.7 RT higher than the one of the IRF-fibril.

**Figure 4.**
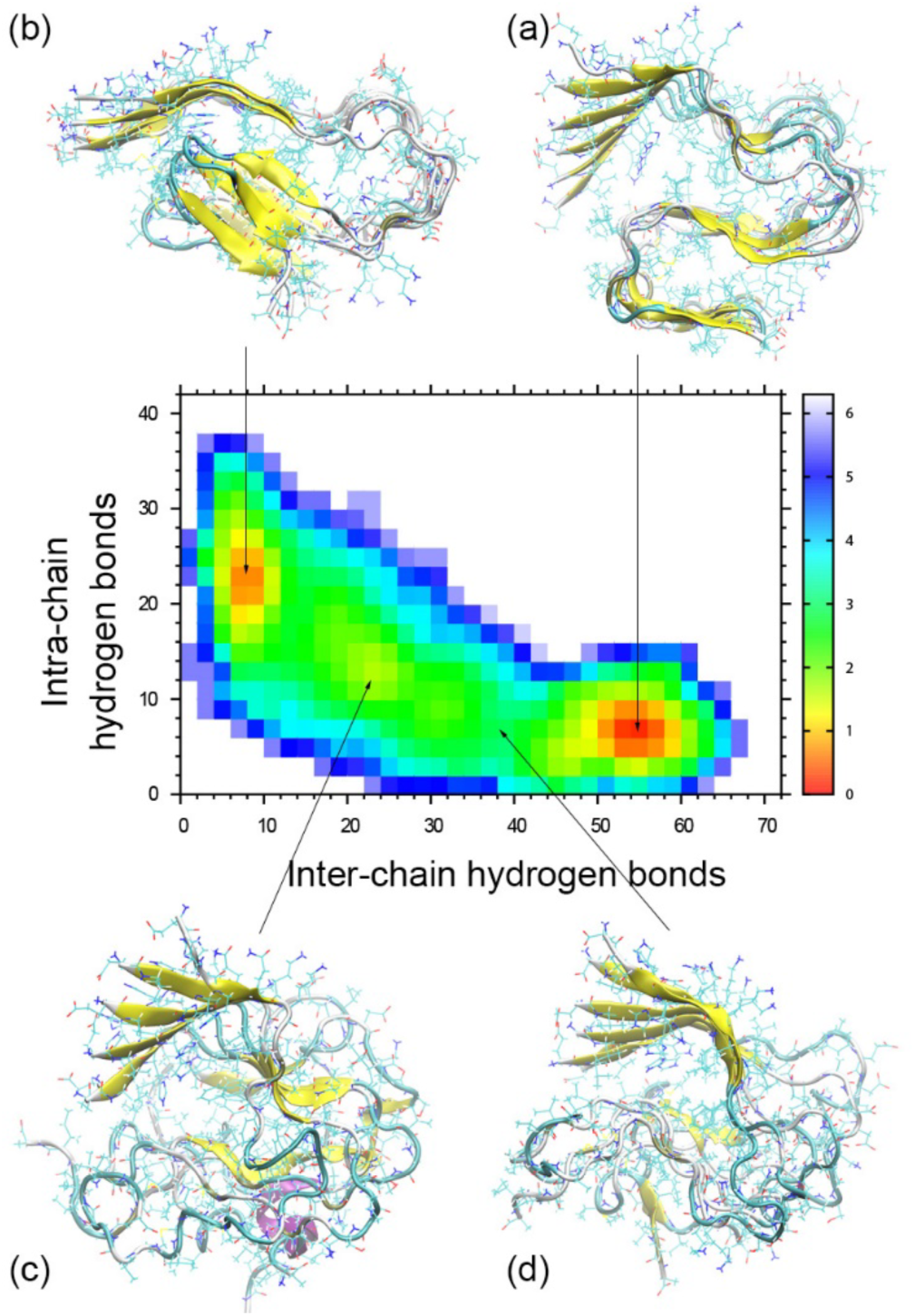
Free energy landscape of the fibril-like hexamer projected on the number of interchain (x-axis) and intra-chain (y-axis) hydrogen bonds involving residues in the C-terminal β2-turn-β3 motif. Data from all 32 replicas are used and reweighted to the case of zero-coupling parameter λ between physical model and biasing Go-model, i.e., the reweighting corrects for the bias introduced by the Go-term. Representative structures are shown for the main regions in the landscape: in-register fibril IRF (a), out-of-register fibril ORF (b), intermediate state (c) and main barrier, the transition state (d).

Given the relative small free energy differences, we conjecture that the ORF fibril organization, while maybe not stable as an assembly with hundreds of peptides, could be a nucleus for growth of fibrils that later fully or partially convert into the more stable in-register form. Such transitions are likely as the barrier between the two forms is only of order a few RT, consistent with the gradual changes observed on the walk shown in Figure 3. Note that the radius of gyration changes little in the landscape if one walks from on fibril arrangement to the other, changing monotonically from 15.8nm to 16.4nm. This suggests that the transition between the two forms does not involve separation of monomers from a fibril that would otherwise change their configuration before re-attaching. Note also that the transition starts from the surface of the fibrils, with the upper layer changing first before deeper layer follow.

### An out-of-register β-barrel model of Aβ42

The Eisenberg group^17^ has proposed that the toxic oligomers in amyloid diseases share a certain (‘cylindrin’) motif made by out-of-register β-strands forming a β-barrel. While it has been shown that eleven-residue long fragments of Aβ peptides (covering residues 25-35) can assemble into this cylindrin form,^18^ steric constraints forbid assemblies of full-sized Aβ chains into this motif if the peptides are in the U-shaped configurations seen in Aβ40 fibrils. This was the motivation for us to assemble trimers and tetramers (constructed with the protocol described in the method section) where the steric constraints are avoided by the S-shaped structure of the individual Aβ_1‒42_ peptides. If stable, our constructs would support the Eisenberg hypothesis that the out-of-register barrel motif is a general architecture for toxic amyloid oligomer.

In Figure 5 (a) and 5(b) we show such trimmers and tetramers, build from residues 27-42, i.e., the out-of-register β2-turn-β3 region of Aβ_1‒42_. Adjacent peptides are distinguished by color. For the tetramer, all backbone hydrogen bonds are in anti-parallel β-sheets formed by the β2-strand and β3-strand of one chain with the corresponding strands of the neighboring chains. The size of the trimer system is comparable with the cylindrin structure developed by Thanh. D et al.^18^ that is made of three chains, each build of two eleven-residue Aβ fragments connected by two glycine (a total of 24 residues), while in our model the relevant part is made of three β-bend fragments consisting of 16 residues (27-42). Note that the size is also similar to that of the cylindrin barrel build from three tandem-repeats of the eleven-residue peptide K11V that also has 24 residues. The cross section of our oligomers is nearly a circle for the trimer, leading to a compact side chain packing of the six strands. For the tetramer the packing is less dense leading to an ellipsoid shape as shown in Figure 5. Both trimer and tetramer have similar height.

**Figure 5.**
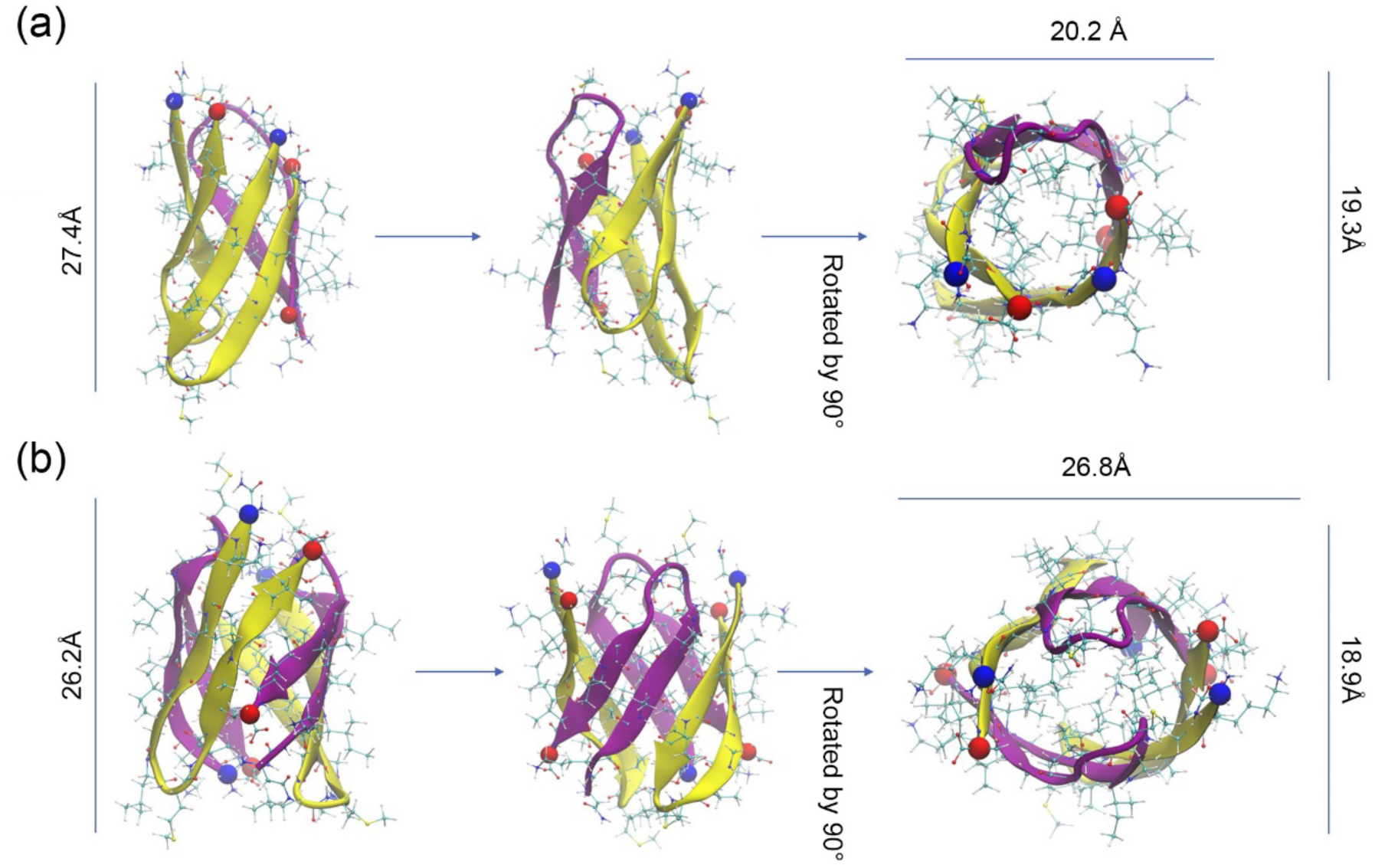
Trimer(a) and tetramer(b) assemblies of Aβ_27‒42_ fragments forming an out-of-register β-barrel. The adjacent peptides are shown in different color. The N-terminal atoms are represented by blue balls, and the C-terminal ones by red balls.

During our molecular dynamic simulations of the Aβ_27‒42_ assemblies the tetramer was more stable than the trimer. The tetramer, shown in Figure 5(b), preserved its β-barrel structure over at least 200ns and changed by only about 2.5 Å while the trimer changed by 2.9 Å. For this reason, we decided to consider only tetramers when building our out-of-register β-barrel assemblies from *full-sized* Aβ_1‒42_ peptides. In the resulting model, the extended β1 sections of adjacent chains form an anti-parallel β-sheet as shown in Figure 6(a), hold together by in-register inter-chain hydrogen bonds. We tried different geometries for the N-terminal residues 1-10, however, in all cases they quickly assumed a random configuration when following the tetramer in molecular dynamic simulations. On the other hand, the tetramer itself is surprisingly stable and preserved its structure over at least 100ns (Figure 6b). This can be seen from Figure 6(c) where we show as function of time the root-mean-square-deviation to the start configuration, evaluated either for the core segment of residues 27-42, for residues 11-42 (i.e., excluding the flexible N-terminal residues), and for the full-size chain. Especially stable is the core region of residue 27-42. The residue 11-26 are constructed as two pairs of anti-parallel sheets, that during the molecular dynamics simulation shrink to residue 12-16, leading to the small root-mean-square deviation seen in Figure 6(d). Overall is the stability of the barrel-shaped tetramer is comparable to the out-of-register fibril model ORF, and likely due to the six hydrogen bonds on the in-register side (between β1 sections of adjacent chains) and four hydrogen bonds in out-of-register side (connecting β2 or β3-regions of one chain with the corresponding strand of a neighboring chain). The side-chains of residues I31, L34 and V40 point inside and form a hydrophobic core for the β-barrel as shown in Figure 6(f). The corresponding binding energies of the core-forming residues 27-42, as approximated by MMGBSA, are shown in Figure 6(e). The distribution of each residue is listed and separated into side-chain and backbone contributions. Note that the side chain of residue I41 does not point inside of the barrel but also contributes to the stability of the barrel by interacting with adjacent chains.

**Figure 6.**
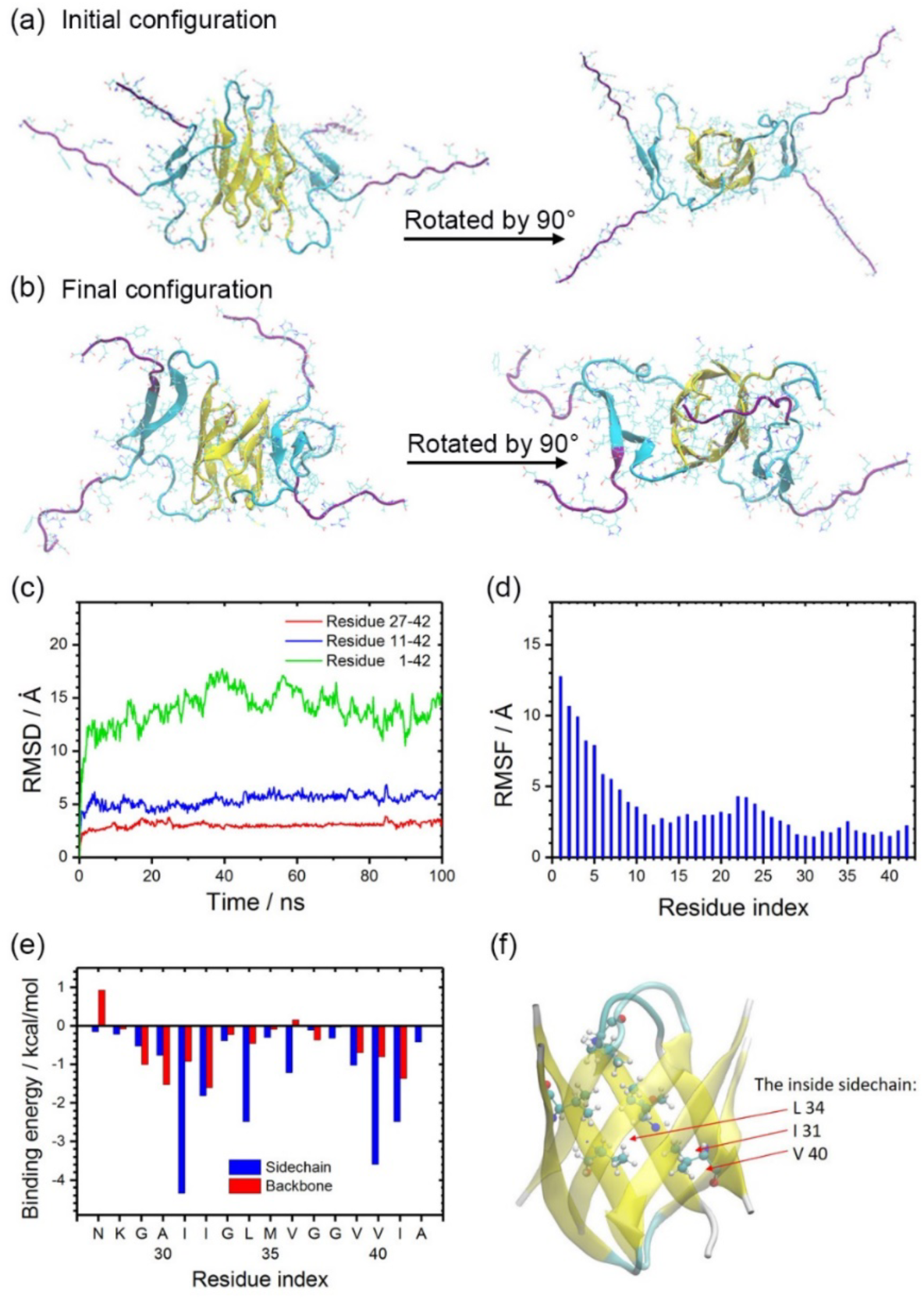
The initial (a) and final (b) configuration of the β-barrel Aβ_1-42_-tetramer. Its root-mean-square deviation (RMSD) to the start configuration as function of time (c) and the root-mean-square fluctuation(RMSF) of all residues(d). Binding energy between chains (c). Shown are only the contributions of residues in the hydrophobic core formed by residues 27-42. Side chain ‐side chain interactions to the binding energy are shown in blue, while the backbone-backbone contributions are drawn in red. (f) Side chain packing inside of the cylindrin-like barrel.

Hence, as seen by us already in an earlier study of cylindrin assemblies^37^ the barrel motif relies on the hydrophobic packing of side chains of residues inside the core, and not on hydrogen bonds between backbone atoms. In our model, the barrel-shaped core is formed by residues 27-42 of the individual chains, i.e., by 16 residues. On the other hand, in earlier work studying K11V-linker^16^ or Aβ_25-35_-linker^18^ models, the cylindrin-like barrel is formed by trimer where each chain has 24 residues. Since the cylindrin barrel are stabilized by the hydrophobic packing of side chains inside the barrel, the smaller number of residues leads to a lower stability of the trimer than seen in the larger-sized chains of the K11V-linker^16^ and Aβ_25-35_-linker^18^ models, and a tetramer is required to provide for sufficient number of stabilizing contacts.

The surprising stability of our tetramer model implies that Aβ_1-42_ peptides can indeed assemble into cylindrin-like barrel with an out-of-register β2-β3 motif. Note that the side chains inside of the barrel are too tightly packed for the barrel o serve as a water channel, which is consistent with previous work simulating barrel-like assemblies of Aβ_25-35_ fragments^18^ and with our measurements. Using an in-house developed plug-in described in our previous work,^42^ we measured an average number of 5.6 water molecules number inside the core region of barrel-like tetramer and 3.3 in the trimer, but did not observe flow of water through the barrel. Hence, while we expect that similar to cylindrin assemblies, our Aβ_1‒42_-barrel are cytotoxic, the toxicity mechanism will likely not be water and ion leakage through cell membranes.

### Out-of-register pathways to Aβ_1-42_ amyloid formation

In previous work^5,43,44^ an in-register pathway of Aβ fibril formation was proposed where monomers assemble into oligomers with in-register U-shaped β-sheets hold together by interchain hydrogen bonds. These oligomers serve as nuclei for the growth of mature fibrils, i.e., are on-pathway to fibril formation. However, structural data on these meta-stable oligomers, ranging in size from dimers to more than 50 chains, are sparse. Especially, there is no direct experimental evidence for the in-register arrangements. On the other hand, our molecular dynamics simulations show that the out-of-register β_2_-turn-β_3_ motif is stable in oligomer and fibrils, which opens the possibility of pathways to fibril formation that utilize the additional degrees of freedom offered by the out-of-register organization.

Hence, while in-register fibrils may form directly by the pathway described above if interchain hydrogen bonding between the β2 and β3 strands of spatially close chains leads to in-register oligomers, we conjecture that these fibrils can also form by another pathway where monomers assemble into small nuclei of out-of-register ORF-like fibril fragments with intrachain hydrogen bonds between β2 and β3 strands. These fragments are metastable and eventually convert into in-register IRF-like assemblies from which the fibril further grows.

Yet another possibility is that Aβ_1-42_ chains with the characteristic out-of-register β2-turn-β3 motif assemble into cylindrin-like β-barrel-shaped oligomers. Such cylindrin-like oligomers have been considered as toxic species in previous work,^18^ and we would expect that our constructs are also cell toxic. Because of their different hydrogen bond organization, these oligomers are separated from the out-of-register fibril ORF-like assemblies by potentially high energetic barriers. However, as both models share the same out-of-register β2-turn-β3 motif, a transition between both organizations may be possible by first dissolving into free monomers that preserve the β2-turn-β3 motif until the chains re-assembled into the other form. In a second step, ORF-like fibrils would then further interconvert into in-register fibrils.

Given the time scale of amyloid formation (hours to days) it is not yet possible to probe the three pathways directly by molecular dynamics. However, both out-of-register pathways rely on the stability of the β_2_-turn-β_3_ motif in *monomers* which can be tested in simulations. We show in Figure 7 results from a 100 ns-long molecular dynamics simulation of the monomer that starts with the peptide in the structure (Figure 7a) it takes in our out-of-register fibril ORF. While the peptide itself is flexible and its final configuration (Figure 7b) after 100ns differs by about 10 Å from the start configuration, the β2-turn-β3 motif is preserved (Figure 7c and d). Note that this is different for S-shaped monomers taken from the in-register fibril model IRF which decay within 10 ns (data not shown) as they are missing the stabilizing hydrogen bonds between the β_2_ and β_3_ strand. Note also that our work is consistent with previous simulations of C-terminal fragments Aβ(1‒42) (x=29‒31,39) that also report a preference for β-hairpin conformation with intra-chain hydrogen bonding.^45^ Interestingly, if residues 41 and 42 are truncated from fragment Aβ_30-42_, the preference shifts to turn-coil states.

**Figure 7.**
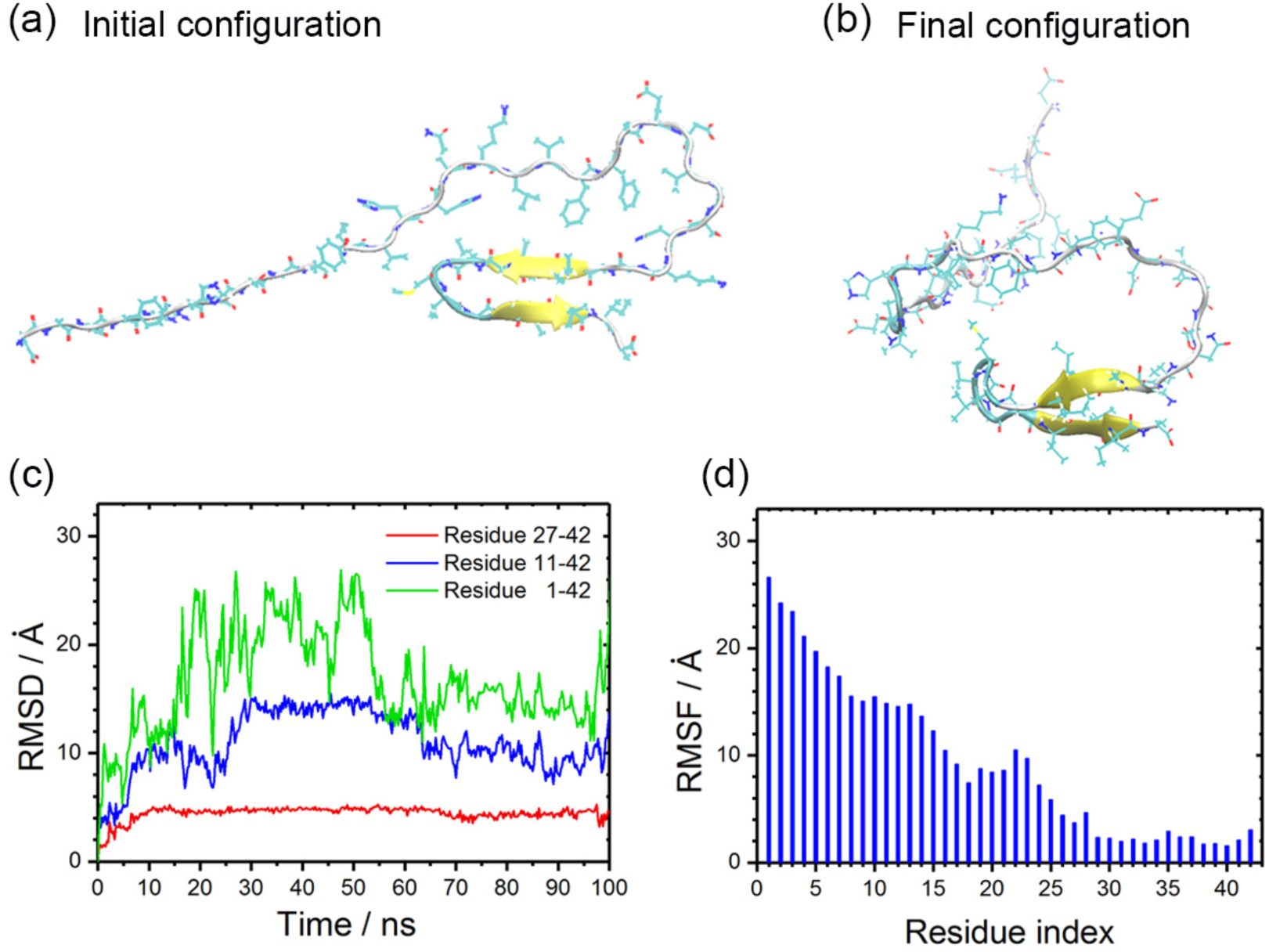
Molecular dynamics simulation of an Aβ_1-42_ monomer. The start configuration (a) has an out-of-register β2-turn-β3 motif hold together by hydrogen bonds between the β_2_ and β_3_ strands. This structure is the same as the peptides assume in the ORF fibril. We show the final configuration of the 100-ns trajectory in (b). The root-mean-square-deviation to the start configuration as function of time is displayed in (c), with the root-mean-square-fluctuations of all residues plotted in (d).

The stability of the β2-turn-β3 motif in Aβ_1‒42_ peptides suggests that the out-of-register pathway leads to higher rate of aggregation as the formation of the critical nucleus will be faster. Once a monomer forms the β_2_-turn-β_3_ motif, it will keep it for considerable time. Time, during which it can interact with others and during which it can form complexes (fibril or barrel-like) that further stabilize it. Formation of such stable structures is much more difficult without the hydrogen-bonding in the β_2_-turn-β_3_ motif.

We remark that while such out-of-register pathways have not yet been detected, their existence could be deduced from signals pointing to high probability of meta-stable intermediate states of monomer with beta-turn motif in residues 27 to 42, or of the K28-A42 salt-bridge which is different from the salt bridge between residues E22 (or D23) and K28 salt-bridge seen in fibrils with U-shaped chains.

## Conclusion

Our guiding assumption is that the higher toxicity of Aβ_1-42_ amyloids over Aβ_1-40_ is due to the ability of Aβ_1-42_ chains to assume three-stranded S-shaped configurations. We have shown that unlike the U-shaped configurations seen for Aβ_1-40_ peptides, S-shaped Aβ_1-42_ chains can form out-of-register antiparallel β-strands arranged as a cylindrical barrel. We have shown that such barrel-shaped tetramers are stable over at last 100ns. These oligomers are likely cytotoxic, as when inserted into cell membranes, they would cause membrane disruption. This by itself may explain the higher toxicity of Aβ_1-42_ peptides over that of the more common Aβ_1-40_ species.

However, there may be a second reason. We could show that the out-of-register β2-turn-β3 motif with its characteristic hydrogen bonding and salt bridge between residues K28 and A42 is even in Aβ_1-42_ monomers surprisingly stable. Monomers with this motif therefore may act as meta-stable intermediates that enhance the chance to form stable oligomers (such as the supposedly cytotoxic barrel structures) and lower the nucleation threshold, which in turn could lead to a higher frequency of toxic species.

The above observation is especially interesting because Aβ_1-42_ peptides can also form fibrils with either in-register or out-of-register hydrogen bonding that are difficult to be distinguished by their chemical shifts in an NMR-experiment. Hybrids of these two fibril arrangements lead to kinks which can explain the “sharp turns” observed in Transmission Electron Microscopy or Atomic Force Microscopy images of Aβ fibrils. We have shown that both arrangements have comparable stability and do interconvert. As the out-of-register fibril arrangement and the barrel-shaped oligomers share the same out-of-register β_2_-turn-β_3_ motif, they likely also interconvert as was seen by us earlier work in simulations of cylindrin assemblies.^37^ We plan to test this hypothesis in future work.

Our results suggest a mechanism where S-shaped monomers, β-barrel oligomer, out-of-register fibrils and in-register fibrils exist in an equilibrium. Its key point is the stability of the out-of-register β_2_-turn-β_3_ motif in Aβ_1-42_ monomers. Such monomers can be formed directly, but can also appear when the S-shaped chains at the end of an in-register fibril convert into out-of-register fibril forms. These out-of-register fibril fragments are less stable and decay to monomers. Once formed, Aβ_1-42_ monomers with the β_2_-turn-β_3_ motif are sufficiently long-lived to either nucleate formation of out-of-register fibrils that in turn may later rearrange into in-register fibrils, or to assemble into the potentially toxic cylindrin-like β-barrel-shaped oligomers. The net-effect of the various transitions and pathways, sketched in Figure 8, is a higher concentration of toxic oligomers than seen for Aβ_1-40_, and a lower effective nucleation threshold.

**Figure 8.**
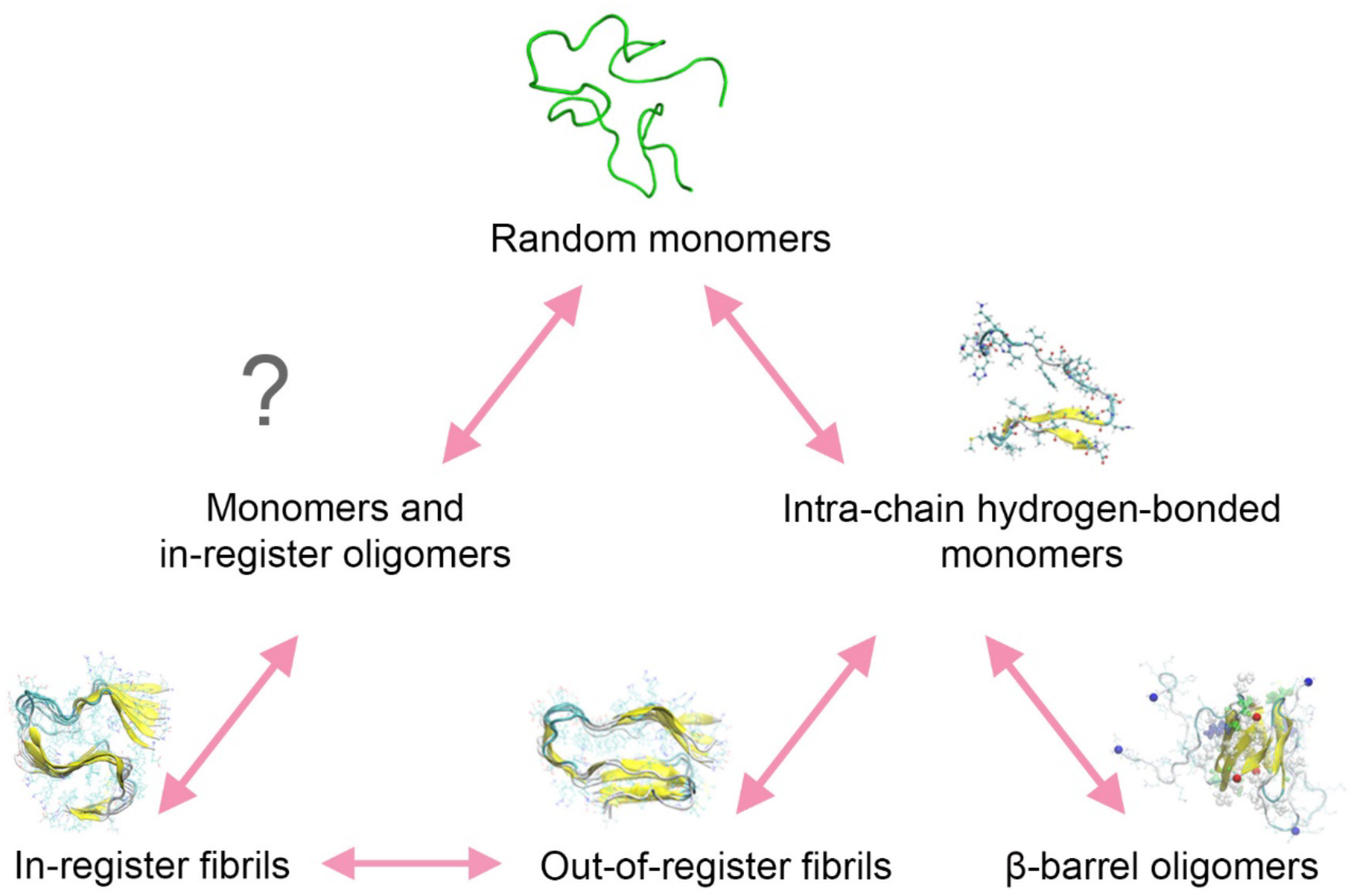
Proposed mechanisms of Aβ_1-42_ amyloid formation relying on the ability of Aβ_1-42_ peptides to assume stable S-shaped configurations.

Probably the most exciting aspect of the proposed mechanism is the possibility that the β_2_-turn-β_3_ motif is “infectious”, i.e., that solvable and therefore mobile monomers (or dimers) with this motif could seed formation of fibril and barrel assemblies by impressing their structure on other monomers. Such self-replicating amyloids are widely discussed^46-53^ and have been observed for the Osaka-mutant^54,55^. We explore now in another ongoing study the possibility of self-replication for S-shaped Aβ_1‒42_ monomers and oligomers.

## Supporting Information

The coordinates of the out-of-register fibril model ORF and of the out-of-register barrel trimer BB3 and tetramer BB4 (made out of either full-sized Aβ_1-42_ chains, or from Aβ_11-42_ or Aβ_27-42_ fragments for BB4) are available in PDB-format as Supporting Information.

## Acknowledgement

The simulations in this work were done using the SCHOONER cluster of the University of Oklahoma and XSEDE resources allocated under grant MCB160005 (National Science Foundation). We acknowledge financial support from the National Institutes of Health under grant GM120578.

